# IEKB: a comprehensive knowledge base for inner ear genetics integrating curated associations, cochlear interactions, Bayesian candidate prioritisation, explainable dark-gene support relations, and a scientific entity network

**DOI:** 10.64898/2026.04.06.716823

**Authors:** Han Wang, Weibang Chen, Hongyu Ning, Yao Cai, Yue Xu, Xuanhe Hou, Lingpin Pang, Ziyin Luo, Chunjie Tian

## Abstract

Inner-ear genetics has expanded rapidly, yet the supporting evidence remains dispersed across a vast literature and across resources that typically emphasise loci, variants, or expression data rather than integrated biological interpretation. Here we present the Inner Ear Knowledge Base (IEKB; https://earkb.org), an open database that unifies curated associations, cochlear interaction evidence, candidate prioritisation, explainable support relations, and network exploration for inner-ear research. IEKB was built with an automated agent-assisted curation workflow that combines schema-constrained literature extraction, continuous human monitoring, and final expert review by inner-ear genetics researchers. By systematically analysing 250,696 PubMed-indexed records retrieved across 16,563 screened genes, IEKB curates 6,051 gene–phenotype–disease associations from 2,494 genes across 43 phenotype categories and 4,102 cochlear gene–gene interactions with pathway, cell-type, and experimental context. IEKB further includes a Bayesian “dark matter” module that prioritises 243,071 candidate gene–phenotype associations for 13,229 genes across all 43 phenotypes (global AUC-ROC = 0.8603; global AUC-PR = 0.1674), together with a supervised dark-relation layer that ranks phenotype-specific known-gene support for each candidate and a multi-entity scientific network containing nearly 4,000 entities, 28,616 deterministic edges, and 83,712 literature-derived relational links. The web resource supports interactive search, multi-parameter filtering, gene-detail pages, bibliometric exploration, domain-specific enrichment against IEKB phenotype and disease gene sets, network visualisation, bulk download in CSV, JSON, SQLite, and XLSX formats, and natural-language evidence-grounded question answering through a companion conversational interface (IEKB QA). To our knowledge, IEKB is the first openly accessible inner-ear resource that integrates curated associations, cochlear interactions, probabilistic candidate prioritisation, auditable known-gene support relations for novel candidates, and a multi-entity scientific network within a single database. All data are released without registration under the CC BY 4.0 license.

## 1 Introduction

Hearing loss (HL) is one of the most prevalent sensory disabilities worldwide, affecting an estimated 1.57 billion individuals—approximately 20% of the global population—according to the Global Burden of Disease Study 2019 [GBD 2019 Hearing Loss Collaborators, 2021]. In industrialized countries, 2–3 out of every 1,000 newborns present with HL, and genetic factors account for approximately 60% of prelingual cases [Morton and Nance, 2006]. Hereditary HL is characterised by extreme genetic heterogeneity: over 170 non-syndromic hearing loss (NSHL) loci have been mapped, implicating more than 124 confirmed genes, and over 400 clinical syndromes include HL as a component [Walls et al., 2026, Shearer et al., 2025]. The advent of next-generation sequencing (NGS) technologies—targeted panels, whole-exome sequencing (WES), and whole-genome sequencing (WGS)—has dramatically accelerated the pace of gene discovery [Cabanillas et al., 2018]. However, the resulting knowledge is dispersed across a vast and rapidly growing literature, making it difficult for researchers and clinicians to assemble an up-to-date and mechanistically informed view of inner ear genetics.

Several publicly available resources partially address this need, yet each captures only one layer of the problem. The Hereditary Hearing Loss Homepage (HHL) provides a valuable catalogue of hereditary hearing loss loci and genes but does not integrate mechanistic context, interaction evidence, or predictive layers [Walls et al., 2026]. The Deafness Variation Database (DVD) offers expert-curated variant classifications for deafness-associated genes, focusing on variant interpretation rather than systems-level organisation [Azaiez et al., 2018]. Gene4HL integrates gene and clinical annotations for hearing-loss-related genes but does not provide cochlear interaction networks, probabilistic candidate prioritisation, or multi-entity network exploration [Huang et al., 2021]. SHIELD provides inner-ear expression resources [Shen et al., 2015], and gEAR hosts community-driven transcriptomic datasets [Orvis et al., 2021]; however, both are centred on expression and do not curate literature-derived phenotype–disease relationships. Consequently, researchers still lack a single, openly accessible platform that connects curated gene–phenotype–disease evidence, cochlear interactions, candidate-gene prioritisation, interpretable support for novel candidates, scientific-network context, and practical data access within one inner-ear resource.

At the same time, recent work in biomedical literature curation has shown that structured extraction can be accelerated by human–AI workflows and interactive curation systems [Sohrab et al., 2022, Wang et al., 2025]. These studies demonstrate the feasibility of semi-automated knowledge structuring, but they are general-purpose tools rather than domain-specific databases for inner ear genetics.

Here we present the Inner Ear Knowledge Base (IEKB), a comprehensive resource for inner-ear genetics built around an automated agent-assisted, expert-curated pipeline. IEKB integrates 6,051 gene– phenotype–disease associations, 4,102 cochlear gene interactions, a Bayesian “dark matter” module prioritising 243,071 candidate associations across 43 phenotypes, a supervised dark-relation layer that provides auditable support rankings from known genes to each dark-matter candidate, a multi-entity scientific network, bibliometric exploration, domain-specific enrichment, downloadable SQLite-backed data access, and a companion natural-language question-answering interface (IEKB QA) that delivers structured, evidence-grounded answers through a multi-phase retrieval-and-synthesis agent pipeline. To our knowledge, IEKB is the first openly accessible inner-ear platform to unify these components within a single resource, providing both broad coverage of the literature landscape and practical tools for hypothesis generation, interpretation, and reuse.

## 2 Materials and Methods

The IEKB construction pipeline consisted of sequential stages encompassing consensus gene-universe construction, prior-knowledge integration, literature retrieval and filtering, agent-assisted curation, post-extraction normalisation, table export, bibliometric analysis, network construction, candidate prioritisation, dark-relation inference, and web deployment. All stages were conducted under researcher monitoring of logs, counts, sampled records, and exported artefacts. Before any output was propagated to the next step or included in the public release, researchers inspected the generated results, and all release-facing biological findings were subsequently reviewed by domain experts in inner-ear genetics. An overview of the pipeline is presented in Figure 1.

**Figure 1:**
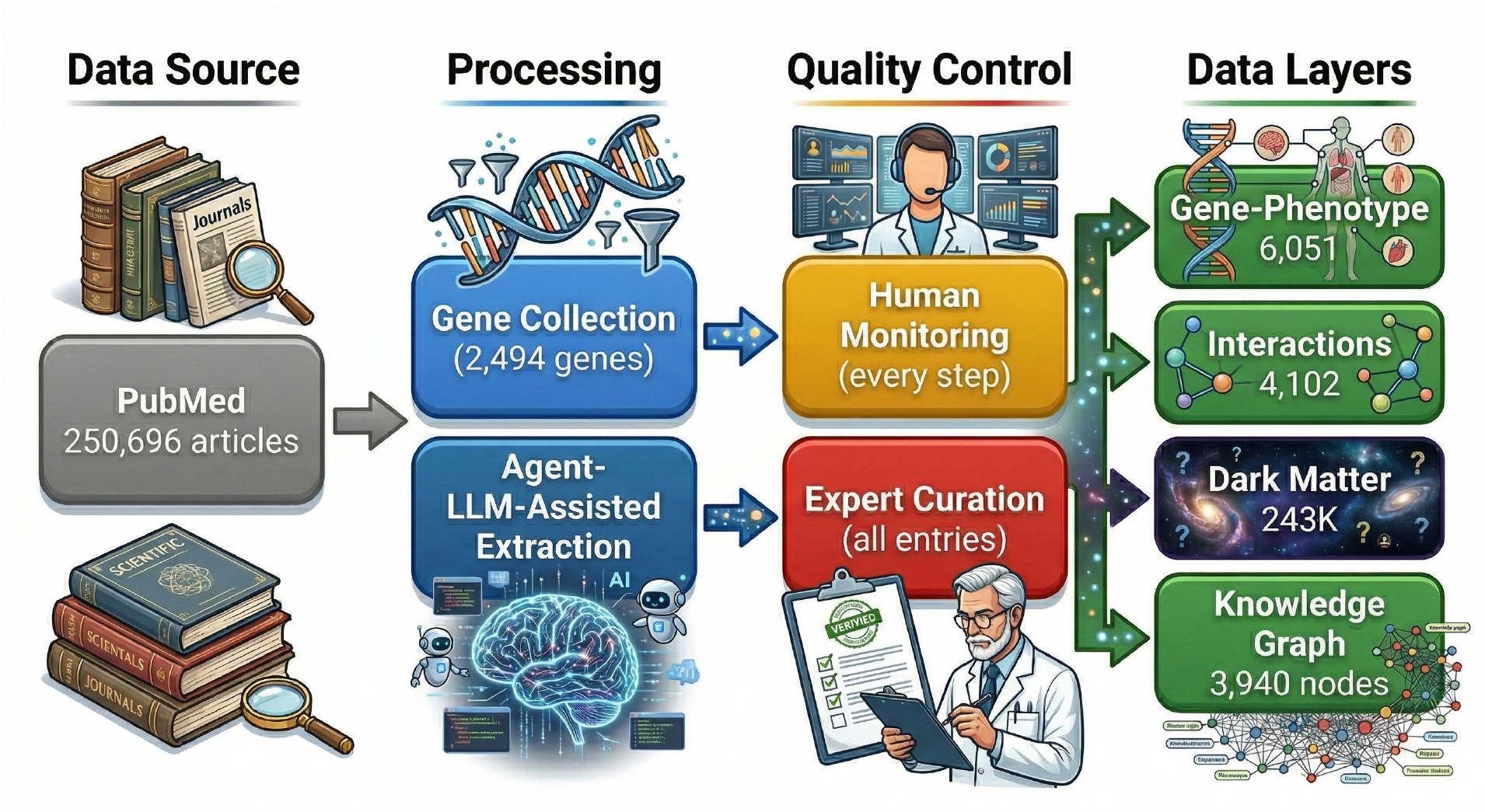
Overview of the IEKB construction pipeline. The workflow proceeds from left to right through four stages. *Data Source*: 250,696 deduplicated PMID-indexed article records were retrieved across 16,563 screened genes by querying PubMed (NCBI Entrez) and Europe PMC with gene-aware inner-ear search strategies. *Processing*: a consensus human gene universe was defined from human protein-coding genes with mouse protein-coding orthologs, followed by two-stage article filtering, agent-assisted structured extraction, and post-extraction normalisation; the current release contains curated evidence for 2,494 genes. *Quality Control*: every computational step was subject to human-in-the-loop monitoring with batch sampling and error correction; all extracted entries were then comprehensively reviewed by domain experts in inner ear genetics. *Data Layers*: the curated pipeline produces five interconnected data layers—6,051 gene–phenotype–disease associations, 4,102 cochlear gene interactions, 243,071 Bayesian dark matter predictions across 43 phenotypes, a supervised dark-relation layer providing auditable known-gene support rankings for dark-matter candidates, and a multi-entity scientific network with nearly 4,000 entities.

### 2.1 Gene universe construction, prior-knowledge integration, and literature retrieval

IEKB first defined a broad human search universe rather than starting from a hand-built hearing-loss seed list. Human and mouse protein-coding genes were downloaded from the NCBI `gene_info` resource, and NCBI `gene_orthologs` was used to retain only human genes with at least one mouse protein-coding ortholog. The resulting consensus human gene universe contained 16,563 genes and was used as the input catalogue for all downstream retrieval steps.

To support downstream prioritisation and network analysis, HGNC metadata were merged for all consensus genes, including approved nomenclature and gene-group annotations [Braschi et al., 2019]. STRING v12 protein annotations and interaction links were then mapped to this universe, and gene–gene interactions were retained when both partners were present in the consensus set and the STRING combined score was at least 400.

Literature retrieval was performed separately for each gene using alias-aware queries built from the approved symbol, recorded synonyms, mouse ortholog symbols, and full gene name. PubMed queries required co-occurrence with 11 inner-ear context terms (“inner ear”, “cochlea”, “cochlear”, “hearing loss”, “deafness”, “hair cell”, “vestibular”, “auditory”, “organ of Corti”, “stria vascularis”, and “spiral ganglion”). PubMed was queried through the NCBI Entrez API [Sayers et al., 2022] and Europe PMC was queried in parallel; each source was capped at 10,000 records per gene, after which results were merged and deduplicated by PMID.

When available, full text was retrieved in priority order from the Europe PMC full-text XML API, NCBI PMC efetch, and Unpaywall-linked open-access landing pages [Piwowar et al., 2018]. Retrieved articles were then passed through a two-stage relevance filter. First, a rule-based prefilter retained only papers whose title or abstract mentioned both the target gene (using the alias set above) and inner-ear-related keywords. Second, an LLM-based classifier (DeepSeek Chat) retained only articles that were both genuinely about the gene and relevant to inner-ear disease biology. Researchers monitored per-gene retrieval counts, sampled matched PMIDs and full texts, and reviewed the resulting gene and article totals before the corpus entered downstream curation. Across the screened gene set, this retrieval layer yielded 250,696 deduplicated PMID-indexed records, and downstream curation produced release-quality phenotype or interaction evidence for 2,494 genes.

### 2.2 Agent-assisted literature curation with human monitoring and expert review

Recent systems such as BiomedCurator and SciDaSynth illustrate how semi-automated workflows can accelerate structured scientific data extraction while preserving opportunities for validation and correction [Sohrab et al., 2022, Wang et al., 2025]. IEKB adapted this paradigm to a domain-specific, per-gene curation workflow. For each gene, an automated literature-synthesis agent processed abstracts and, when available, full text in multi-pass evidence batches and produced two categories of structured output: (i) gene–phenotype–disease associations and (ii) gene–gene interactions in the cochlea. Outputs were constrained by Pydantic data schemas [Colvin, 2024]. The gene–phenotype schema captured identifiers, phenotype and disease labels, inheritance, mechanistic interpretation, cochlear and cellular context, model-system evidence, supporting PMIDs, and a curator-facing summary. The interaction schema captured gene pairs together with interaction type, directionality, regulatory effect, pathway and biological context, cell type, experimental method, evidence level, confidence, and supporting PMIDs. Post-extraction, per-gene outputs were passed through a hybrid cleaning and alignment stage that combined HGNC symbol standardisation [Braschi et al., 2019], rule-based normalisation, and LLM-assisted vocabulary mapping across phenotype, cell-type, cochlear-region, pathway, and biological-context fields. Duplicate entries were merged, evidence from multiple sources was consolidated, and empty placeholder records were generated for genes missing from the raw extraction output so that a complete per-gene index could be exported. Researchers inspected the cleaned per-gene files and exported tables before release-facing results were frozen.

Human-in-the-loop monitoring was applied throughout extraction, cleaning, and export. Researchers randomly sampled approximately 10% of each extraction batch and cross-referenced agent outputs against the source literature, flagging and correcting errors. Entries that failed schema validation, exhibited low confidence scores, or contradicted prior knowledge were automatically flagged for manual review. Systematic error patterns identified during sampling were fed back into the extraction prompts and validation rules to iteratively improve output quality. All 6,051 gene–phenotype–disease associations and 4,102 gene–interaction entries in the public release were subsequently reviewed by domain experts in inner ear genetics, who confirmed or corrected each record’s evidence level, mechanism description, and inheritance pattern by cross-referencing with ClinGen [DiStefano et al., 2019] and HHL [Walls et al., 2026]. Disagreements between annotators were resolved through multi-expert consensus discussion.

### 2.3 Quality assessment and confidence classification

To quantify curation quality, a random sample of automatically extracted records was independently evaluated by two annotators in a double-blind design. Precision and recall were computed against an expert-curated gold standard, and inter-annotator agreement was measured using Cohen’s kappa. All records were assigned to one of two confidence tiers: Tier 1 entries are explicitly supported by the source literature and confirmed by expert review; Tier 2 entries are inferred from indirect evidence and flagged by experts as requiring further validation. Researchers reviewed the resulting metric tables and discrepant cases before the final quality summaries were incorporated into the release manuscript and database. Detailed quality metrics are reported in Table 1.

**Table 1:**
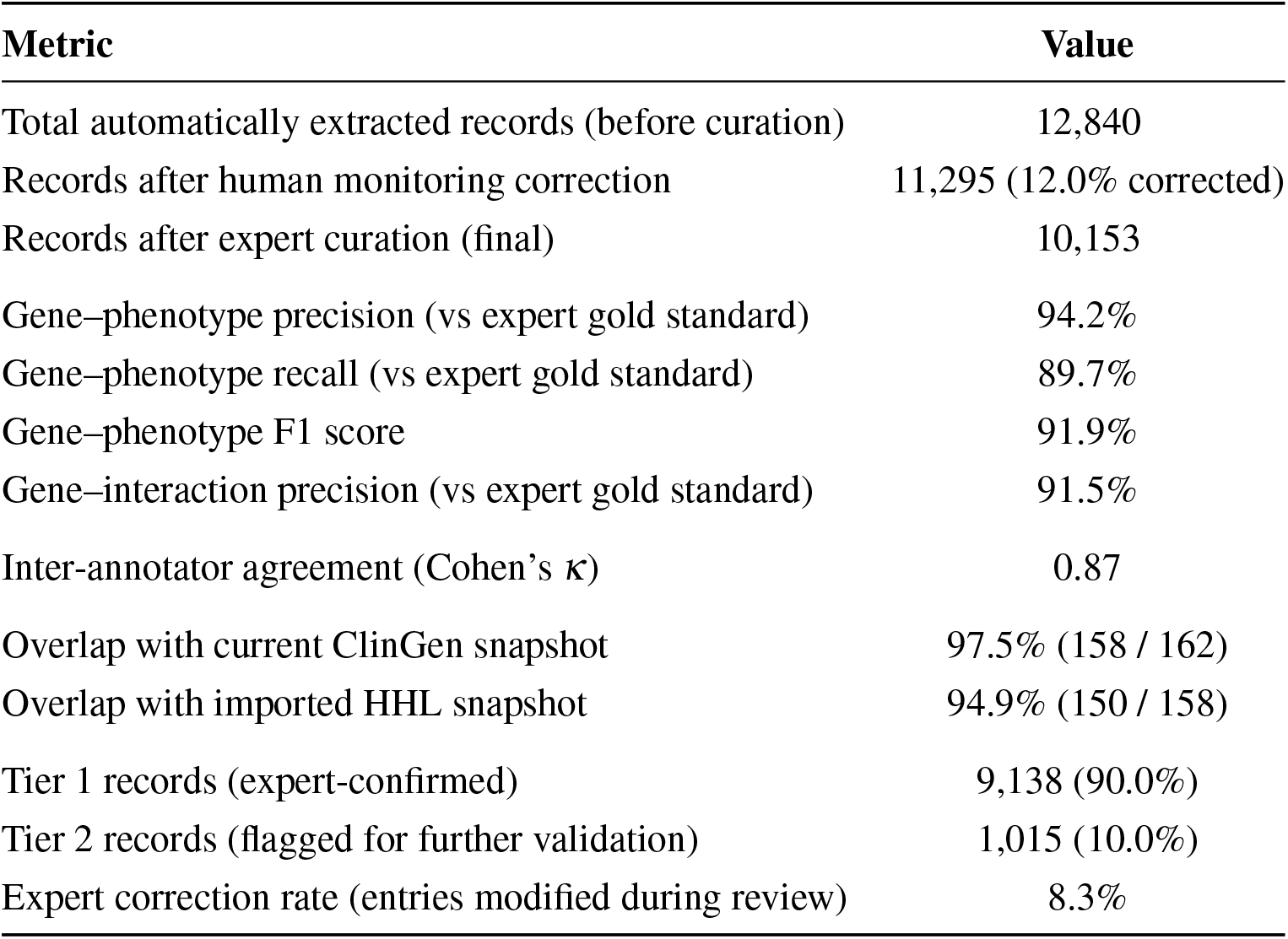
Quality assessment of automated extraction and expert curation. Metrics were computed on a randomly sampled subset of records evaluated independently by two annotators. Precision, recall, and F1 are reported against the expert-curated gold standard. Tier 1 entries are explicitly supported by literature and confirmed by experts; Tier 2 entries are inferred from indirect evidence and flagged for further validation.

| Metric | Value |
| --- | --- |
| Total automatically extracted records (before curation) | 12,840 |
| Records after human monitoring correction | 11,295 (12.0% corrected) |
| Records after expert curation (final) | 10,153 |
| Gene–phenotype precision (vs expert gold standard) | 94.2% |
| Gene–phenotype recall (vs expert gold standard) | 89.7% |
| Gene–phenotype F1 score | 91.9% |
| Gene–interaction precision (vs expert gold standard) | 91.5% |
| Inter-annotator agreement (Cohen’s $\kappa$) | 0.87 |
| Overlap with current ClinGen snapshot | 97.5% (158 / 162) |
| Overlap with imported HHL snapshot | 94.9% (150 / 158) |
| Tier 1 records (expert-confirmed) | 9,138 (90.0%) |
| Tier 2 records (flagged for further validation) | 1,015 (10.0%) |
| Expert correction rate (entries modified during review) | 8.3% |

### 2.4 Bayesian dark matter inference

To prioritise novel gene–phenotype associations beyond those already curated from the literature, IEKB implements a probabilistic “dark matter” framework informed by network-propagation ideas widely used in disease-gene discovery [Tong et al., 2008, Visonà et al., 2024]. Inputs comprised the curated gene– phenotype table, the STRING-derived interaction graph, and HGNC gene-group annotations. Phenotype labels were first normalised by merging exact duplicates and collapsing sparse categories according to a predefined mapping, yielding 43 analysis categories in the current release. Genes with at least one Step 2.5-filtered article were treated as seed genes; genes with raw retrieved literature but zero filtered articles were treated as reliable negatives; the remaining genes were scored as dark-matter candidates.

Let *S*_*p*_ denote the seed-gene set for phenotype *p, A* the weighted STRING adjacency matrix, *d*_*g*_ the degree of gene *g*, and *G*_*g*_ the HGNC gene-group set of gene *g*. For each seed gene *j* ∈ *S*_*p*_, a quality weight was defined as

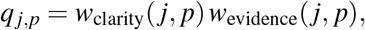

where *w*_clarity_ ∈ {1.0, 0.6, 0.3} for clear, potential, and unknown mechanism assignments, respectively, and *w*_evidence_ ∈ {1.0, 0.9, 0.8, 0.5, 0.4} for *in vivo, clinical, in vitro, review*, and *computational* evidence.For each phenotype *p* and gene *g*, we computed a three-feature vector *x*_*g,p*_ = (*f*_1_, *f*_2_, *f*_3_). The first feature was a direct-neighbour score on the STRING graph,

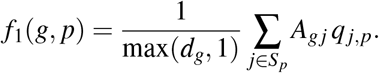

The second feature was a Random Walk with Restart (RWR) score on the row-normalised transition matrix *W*,

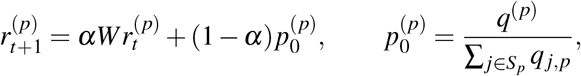

with *α* = 0.7 by default and 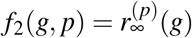 at convergence. The third feature quantified overlap with phenotype-linked gene groups,

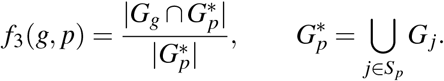

For each phenotype, a class-balanced logistic-regression model transformed these features into a raw score,

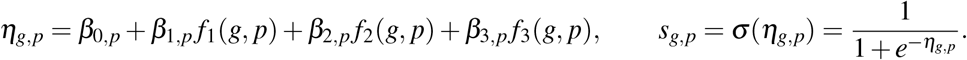

Out-of-fold scores from cross-validation were calibrated globally by Platt scaling,

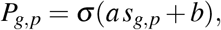

which yielded the posterior-like prioritisation score used in the release. Model performance was evaluated independently for each phenotype using stratified *k*-fold cross-validation over seed genes and reliable negatives, with five folds when at least 25 seed genes were available and three folds otherwise. Final phenotype-specific models were then trained on all available seed and reliable-negative genes, applied to dark-matter genes, and converted to calibrated scores through the learned scaler. Within each phenotype, Benjamini–Hochberg correction [Benjamini and Hochberg, 1995] was applied to 1 − *P*_*g,p*_ to obtain *q*-values. Researchers monitored phenotype merges, feature distributions, cross-validation outputs, and ranked prediction tables at every run; all release-facing validation summaries were checked by researchers, and top-ranked novel candidates were reviewed by domain experts for biological plausibility.

### 2.5 Dark relation inference

While the Bayesian dark matter module assigns a posterior score to each candidate gene–phenotype pair, it does not explain *which* known genes most strongly support a given prediction. To provide auditable evidence chains, IEKB adds a supervised dark-relation layer that ranks known genes as phenotype-specific supporting evidence for each dark-matter candidate.

#### Problem formulation

The ranking task is query-centric. During training, each query consisted of a phenotype *p* and a held-out target gene *g*^⋆^ drawn from the curated seed set for that phenotype. Candidate source genes were labelled positive if they were other known genes associated with the same phenotype and negative if they were known genes associated with different phenotypes. At inference time, the held-out target was replaced by a dark-matter candidate gene, yielding the same ranking pattern exposed in the web interface: for each (dark gene, phenotype) pair, which known genes provide the strongest support.

#### Feature engineering

For each source–target gene pair (*j, g*^⋆^), a ten-dimensional feature vector was computed:

1. *Direct network score*: the STRING edge weight *A*_*j,g*_⋆ .
2. *RWR score*: the Random Walk with Restart stationary probability from *j* to *g*^⋆^ on the row-normalised STRING matrix (*α* = 0.7).
3. *Gene-group overlap*: the Jaccard-like ratio |*G*_*j*_ ∩ *G*_*g*_⋆ |/|*G*_*j*_ ∪ *G*_*g*_⋆ | of HGNC gene-group memberships.
4. *Shared-neighbour ratio*: |𝒩(*j*) ∩ 𝒩(*g*^⋆^)|/|𝒩(*j*) ∪ 𝒩(*g*^⋆^)|.
5. *Adamic–Adar index*: ∑_*w*∈𝒩(*j*)∩𝒩(*g*_⋆) 1/ln *d*_*w*_, a link-prediction measure that up-weights shared neighbours of low degree [Adamic and Adar, 2003].
6. *Direct–RWR interaction*: the product of features 1 and 2.
7. *Source log-degree*: log(1 + *d* _*j*_).
8. *Target log-degree*: log(1 + *d*_*g*_⋆).
9. *Source log-evidence*: log(1 + *e* _*j*_), where *e* _*j*_ is the aggregated evidence weight for the source gene across the curated corpus.
10. *Target log-evidence*: log(1 + *e*_*g*_⋆), defined analogously for the target gene.

#### Model training and selection

Training pairs were assembled across all qualifying phenotypes (≥ 6 seed genes) using hard-negative mining, with 50% of negatives chosen as phenotype-confusable genes showing the strongest direct network or gene-group proximity to the target. A class-balanced logistic regression served as the default model; when available, a LightGBM gradient-boosted classifier was trained using the same query grouping. Both models were evaluated with grouped *k*-fold cross-validation (*k* = 5), ensuring that all pairs sharing a query appeared in the same fold. Macro-averaged Recall@10 and AUPR were then computed over held-out queries. Model selection between logistic regression and LightGBM was based on a paired bootstrap comparison (*B* = 400 iterations, 95% confidence): LightGBM was preferred only if it showed a bootstrap-supported improvement in macro Recall@10 without materially degrading calibration (Brier score increase ≤ 0.02, macro AUPR decrease < 0.005).

#### Calibration and inference

Out-of-fold raw scores were calibrated via Platt scaling. The winning model was then retrained on the full training set and applied to all dark-matter candidates whose posterior exceeded a configurable threshold (default 0.1). For each (dark gene, phenotype) pair, known genes were ranked by calibrated relation score; the top *K* relations (default *K* = 10, minimum score ≥ 0.05) were retained for export.

#### Explainability

Each exported relation includes the calibrated score, the individual feature values, and a natural-language evidence summary highlighting the three strongest contributing signals (e.g., direct STRING support, network diffusion, or shared gene-group membership), thereby enabling researchers to audit why a particular known gene was surfaced.

### 2.6 Knowledge graph construction

A multi-entity scientific network was constructed to represent the relational landscape of inner ear biology, following the broader biomedical knowledge-graph paradigm of integrating curated resources with literature-derived relations [Nicholson and Greene, 2020]. The first layer is a deterministic network builder that expands the structured columns of the curated phenotype/disease and interaction tables into six core entity categories: gene, phenotype, disease, pathway, cell type, and cochlear region. This layer generates gene–phenotype, gene–disease, gene–cell-type, gene–region, gene–gene, and gene–pathway edges.

The second layer applies an LLM-based link extractor to each curated finding and emits additional directed semantic links with normalised entity names, relationship verbs, and low/medium/high strength labels. This literature-derived layer can introduce broader entity types such as molecules and biological processes beyond the six core deterministic categories. Evidence strength in the deterministic layer was assigned from supporting PMID counts and, for curated interactions, available confidence labels. Researchers reviewed node and edge counts and sampled deterministic and LLM-derived links after each build. Disease-specific and cell-type-specific sub-networks were generated for web visualisation, and key sub-networks—including those for Usher syndrome and DFNB1—were reviewed by experts for biological plausibility.

### 2.7 Bibliometric analysis

Publication metadata for all 250,696 deduplicated PMID-indexed articles retrieved before Step 2.5 filtering were analysed to characterise temporal trends, journal distributions, citation patterns, and keyword frequencies. Summary statistics (year range 1814–2026, 8,285 unique journals, 4,488,908 total citations, mean 17.91 citations per article, and 14.4% open access rate) were computed and made available through the web interface; the interactive trend plots focus on the post-1970 period for readability. All bibliometric results were reviewed by researchers to confirm the absence of data-processing artefacts before the release summaries were finalised.

### 2.8 Database design and web implementation

The web build step compiled the exported artefacts into a SQLite database with 11 tables covering curated associations, curated interactions, dark-matter predictions and validation results, dark-relation support rankings, network nodes, deterministic edges, literature-derived links, article metadata, and imported ClinGen and HHL reference snapshots. The web application was built with Next.js 16 and employs sql.js to execute SQL queries entirely within the user’s browser, eliminating the need for a backend server. The front end uses Cytoscape.js for interactive network visualisation [Franz et al., 2016], Recharts for statistical charts, TanStack Table for paginated data tables, and Fuse.js for client-side fuzzy search. The application is deployed on Cloudflare Pages with large data files hosted on Cloudflare R2. After data import, researchers performed end-to-end functional testing of the web interface to verify that all displayed data matched the source CSV files, and the public biological content surfaced by the application inherited the expert-reviewed release tables described above.

### 2.9 Evidence-grounded question answering (IEKB QA)

To complement the browsing-oriented web interface, IEKB provides a companion natural-language question-answering service, IEKB QA (https://qa.earkb.org), that allows researchers to pose complex biomedical questions in free text and receive structured, citation-backed answers grounded in the IEKB database.

The system operates through a multi-phase agent pipeline. First, incoming questions are classified by intent (e.g. gene lookup, disease query, target discovery, statistical inquiry) and biomedical entities such as gene symbols and disease terms are extracted. When the question is complex, an optional planning stage decomposes it into a sequence of retrieval sub-tasks. Each sub-task issues schema-constrained SQL queries against the same IEKB SQLite data source used by the main web application, retrieving gene– phenotype associations, cochlear interactions, dark-matter predictions, dark-relation support rankings, network edges, and supporting literature as needed. Users may choose to include or exclude dark-matter and dark-relation evidence at query time, matching the toggle provided on the main site.

Retrieved evidence is then passed through a deterministic candidate-ranking stage that scores genes across multiple dimensions—including disease association strength, mechanism clarity, evidence level, interaction connectivity, and dark-matter posterior probability—before a large language model synthesises the ranked evidence into a coherent, citation-backed narrative answer. A post-synthesis re-ranking step aligns the final candidate list with the LLM-generated text to ensure consistency between the narrative and the structured output. The entire pipeline is streamed to the browser via Server-Sent Events (SSE), so users can observe each reasoning phase in real time. IEKB QA is compatible with any OpenAI-compatible language-model API and does not depend on a specific model provider. The conversational interface supports multi-turn follow-up questions within the same evidence session, and completed sessions can be exported for record-keeping.

## 3 Database Content and Usage

All data in IEKB were produced through the agent-assisted, human-monitored pipeline described above, with every public entry reviewed before release.

### 3.1 Quality assessment results

Prior to large-scale release, we evaluated the accuracy of the automated extraction workflow on a randomly sampled subset of records. Cross-validation against expert-curated gold standards demonstrated high precision for both gene–phenotype associations and gene–gene interactions (Table 1). Human-in-the-loop monitoring identified and corrected recurrent issues including unsupported mechanism summaries, entity normalisation mismatches, and imprecise evidence assignment. After comprehensive expert review, the final release comprises predominantly Tier 1 records (entries explicitly supported by the literature and confirmed by experts), with a smaller Tier 2 subset flagged for further validation.

### 3.2 Overview of IEKB

The current release of IEKB contains 6,051 curated gene–phenotype–disease associations derived from 2,494 unique human genes across 43 phenotype categories (Figure 2). The most frequently represented phenotype is sensorineural hearing loss (964 associations), followed by inner ear malformation (480), noise-induced hearing loss (364), vestibular dysfunction (343), and age-related hearing loss/presbycusis (335). Gene–phenotype entries are annotated with inheritance patterns (autosomal recessive, autosomal dominant, X-linked, mitochondrial, digenic, or complex), mechanism clarity ratings (clear, potential, or unknown), affected cell types, cochlear regions, animal model information, and supporting PubMed identifiers. Evidence annotations span *in vivo, clinical, in vitro, review*, and *computational* support, often in combination, with experimental evidence predominating in the curated corpus.

**Figure 2:**
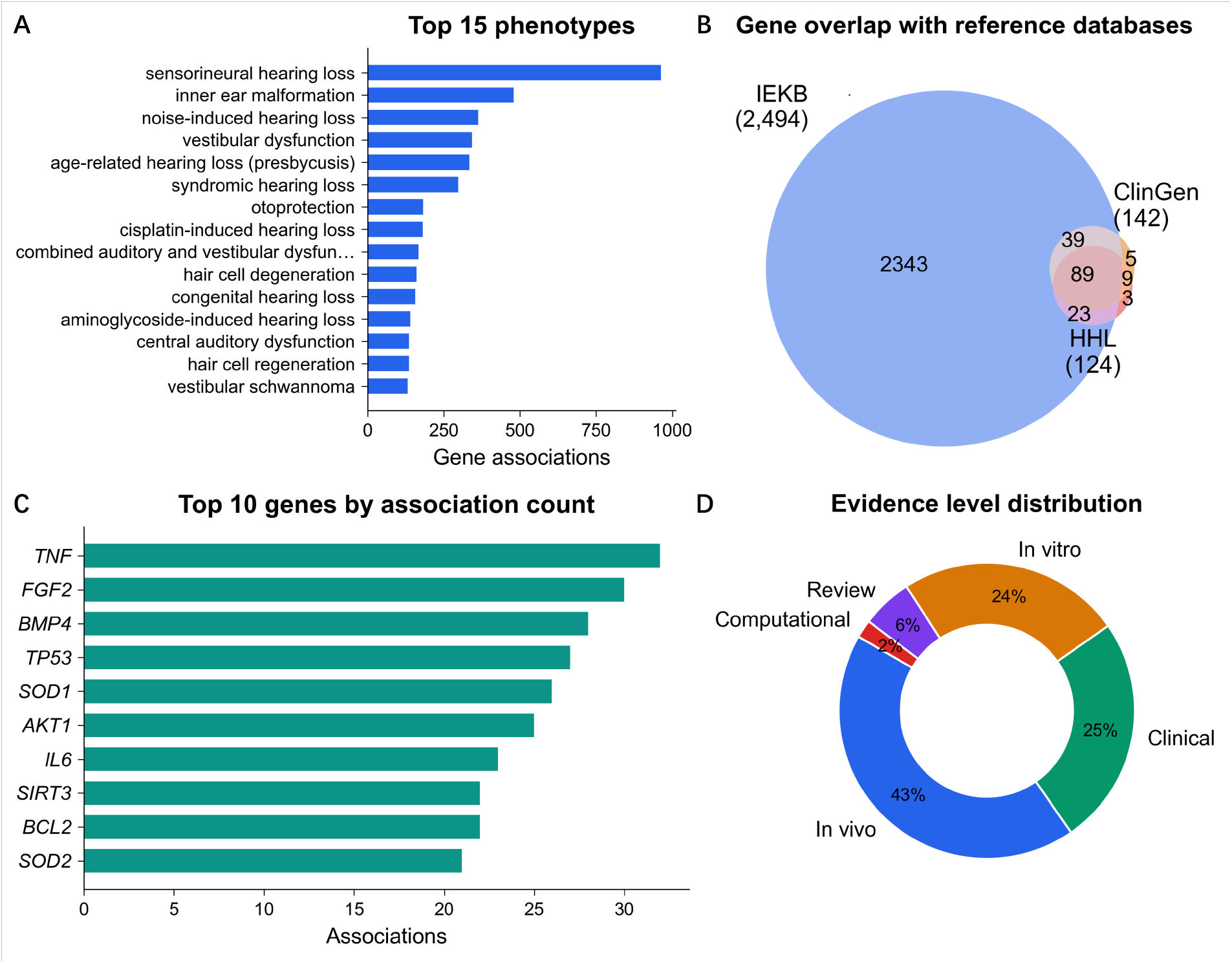
Database content overview. **(A)** Distribution of gene–phenotype associations across the top inner-ear phenotype categories, with sensorineural hearing loss, inner ear malformation, and noise-induced hearing loss among the most represented terms. **(B)** Venn diagram illustrating the overlap between IEKB genes and the imported ClinGen and HHL reference sets, highlighting both strong coverage of established hearing-loss genes and substantial expansion beyond existing catalogues. **(C)** Top 10 genes ranked by the number of phenotype associations. *TNF, FGF2*, and *BMP4* are among the most connected genes, reflecting both established hearing-loss genes and pleiotropic regulators implicated in otoprotection. **(D)** Distribution of evidence annotations across all 6,051 gene–phenotype entries, showing that experimental and clinical evidence dominate the current release.

Comparison with established reference databases demonstrates both high coverage and complementary breadth. In the current imported reference snapshot, 158 of 162 ClinGen hearing-loss genes (97.5%) and 150 of 158 HHL entries (94.9%) are represented in IEKB. Conversely, 2,301 IEKB genes are absent from both reference sets, reflecting substantial expansion beyond established hearing-loss catalogues.

Highly connected genes by phenotype-association count include *TNF* (32), *FGF2* (30), *BMP4* (28), *TP53* (27), and *SOD1* (26), reflecting both established hearing-loss genes and pleiotropic regulators implicated in otoprotection and cochlear stress responses.

### 3.3 Gene–gene interactions

IEKB catalogues 4,102 cochlear gene–gene interactions, each annotated with interaction type, directionality, regulatory effect, pathway context, biological context, cell type, experimental methods, evidence level, species, and confidence rating. The most prevalent interaction types are signalling pathway interactions (1,433), transcriptional regulation (875), and protein–protein interactions (731), consistent with the importance of developmental signalling pathways such as Notch, Wnt, and FGF in inner ear development and patterning [Nakajima, 2015].

### 3.4 Bayesian dark matter prioritisation

The dark matter inference model generated 243,071 predictions spanning 13,229 candidate genes across all 43 phenotype categories (Figure 3). Cross-validation yielded a global AUC-ROC of 0.8603 and a global AUC-PR of 0.1674. Per-phenotype AUC-ROC values ranged from 0.6981 (sensorineural hearing loss, the broadest category with 786 seed genes) to 0.9758 (infection-induced hearing loss, 38 seeds), with 42 of 43 phenotypes classified as “reliable” and one (sensorineural hearing loss) classified as “low reliability” owing to the heterogeneity of its seed-gene set. The separation between posterior distributions for known seeds and novel predictions indicates that the model effectively distinguishes established associations from previously unsurfaced candidates. As a case study, we examined the top 15 novel candidates for sensorineural hearing loss. The highest-ranked gene, *RBM5* (posterior = 0.81), is therefore highlighted as a biologically plausible candidate for further investigation.

**Figure 3:**
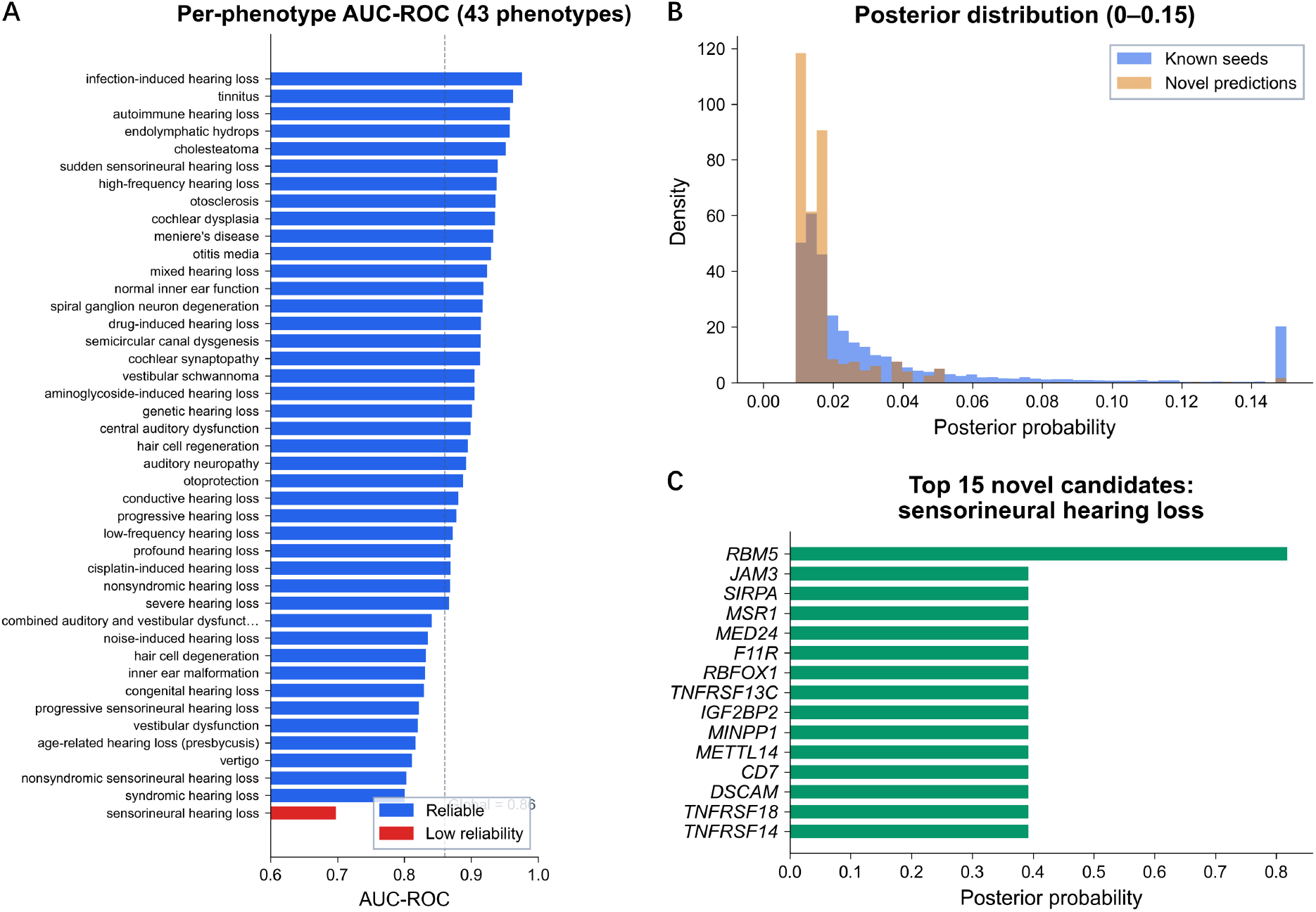
Bayesian dark matter predictions. **(A)** Per-phenotype AUC-ROC for the Bayesian dark matter inference model across all 43 phenotype categories, sorted in descending order. Blue bars indicate phenotypes with “reliable” cross-validation metrics; the single red bar indicates “low reliability” (sensorineural hearing loss, AUC-ROC = 0.698, 786 seed genes). The dashed line marks the global AUC-ROC of 0.86. Values range from 0.698 to 0.976 (infection-induced hearing loss, 38 seeds). **(B)** Posterior probability density distributions (range 0–0.15) for known seed genes (blue) versus novel predictions (amber). Known seeds exhibit a broader distribution extending to higher posterior values, while novel predictions are concentrated near zero, confirming that the model discriminates between established and candidate associations. **(C)** Top 15 novel candidate genes for sensorineural hearing loss ranked by posterior probability. *RBM5* (posterior = 0.81) is the highest-ranked novel candidate. All listed genes lack retained filtered literature evidence for sensorineural hearing loss and are prioritised through local network-neighbour, network-propagation, and gene-group features. The dark-relation layer further provides auditable known-gene support rankings for each candidate, showing which established hearing-loss genes contribute the strongest phenotype-specific evidence.

### 3.5 Dark relation support rankings

To make the dark matter predictions actionable, the dark-relation layer provides an auditable evidence chain linking each novel candidate to established hearing-loss genes. For every (dark gene, phenotype) pair passing the posterior threshold, the model ranked all known genes associated with that phenotype by calibrated relation score. Model selection via paired bootstrap testing identified the release model on the basis of grouped cross-validation performance, after which the winning classifier was retrained on the full training set and applied to all qualifying dark-matter candidates.

Each exported relation includes the calibrated score, the underlying feature decomposition (direct STRING edge weight, RWR network-diffusion score, gene-group overlap, shared-neighbour ratio, and Adamic–Adar index), and a natural-language evidence summary highlighting the three strongest contributing signals. This design allows researchers to inspect not only *which* known genes support a dark-matter prediction but also *why*, thereby facilitating more informed experimental follow-up.

Within the web interface, these rankings convert a posterior score into an inspectable support profile: users can examine how support is distributed across phenotypes, compare the strongest known-gene supporters for a candidate, and trace the network-derived features that drive each ranking.

### 3.6 Scientific entity network

The IEKB scientific network links genes, diseases, phenotypes, pathways, cell types, and cochlear regions through 28,616 deterministic edges and 83,712 literature-derived relational links, yielding a graph of nearly 4,000 connected entities (Figure 4). The degree distribution follows an approximate power-law, characteristic of biological networks. The highest-degree hub entities are organ of Corti (degree 1,188), hair cells (871), sensorineural hearing loss (763), spiral ganglion neurons (743), and spiral ganglion (656), reflecting the centrality of these structures and phenotypes in inner ear biology. Among pathway hubs, Oxidative Stress Response (degree 135), Actin Cytoskeleton Regulation (125), Developmental Signaling (123), FGF Signaling (116), and PI3K/Akt Signaling (107) emerged as the most connected, highlighting the joint prominence of developmental and stress-response programmes in the IEKB network.

**Figure 4:**
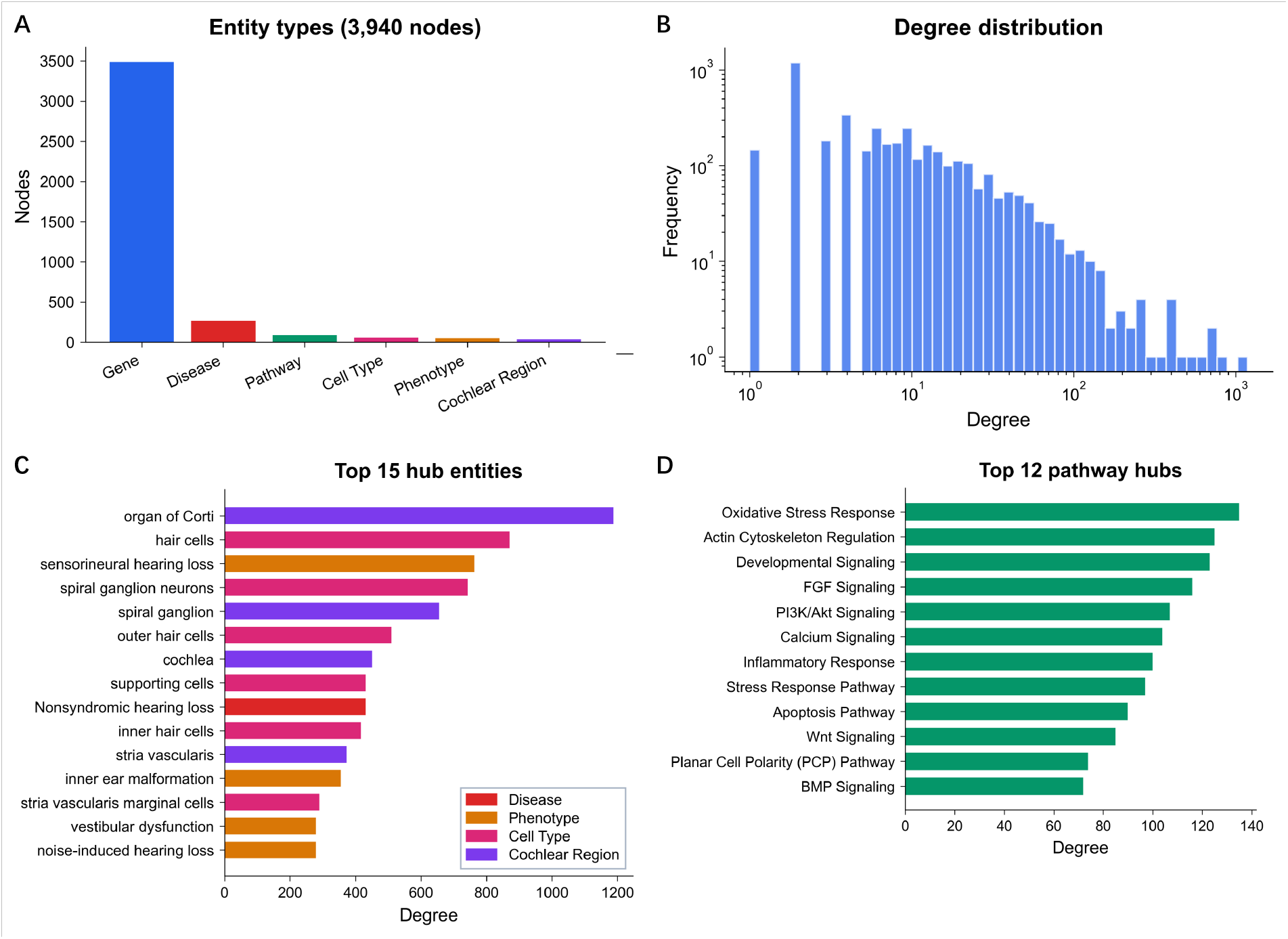
Scientific entity network analysis. **(A)** Distribution of entity types across the IEKB scientific network. **(B)** Node degree distribution plotted on log–log axes, exhibiting an approximate power-law shape characteristic of scale-free biological networks. **(C)** Top hub entities ranked by degree, highlighting the central positions of organ of Corti, hair cells, and sensorineural hearing loss within the network. **(D)** Top pathway hubs ranked by degree, with Oxidative Stress Response, Actin Cytoskeleton Regulation, and Developmental Signaling among the most connected pathways.

### 3.7 Case applications

To demonstrate the utility of IEKB, we present two representative use cases. An example of the gene detail page is shown in Figure 6 for *ABCC1*.

#### Case 1: GJB2 and DFNB1

*GJB2* (connexin 26) is the most commonly mutated gene in non-syndromic hearing loss, responsible for the DFNB1 locus [Kelsell et al., 1997]. Querying IEKB for *GJB2* retrieves all associated phenotypes, interaction partners, dark matter co-predictions, and, where available, dark-relation support links showing which known genes most strongly support nearby dark-matter candidates. The interaction network reveals connections to *GJB6, GJB3*, and *GJA1* through gap junction complex formation, as well as links to the Wnt and Notch signalling pathways. The enrichment analysis module enables users to test whether a set of DFNB1-associated genes is enriched for specific IEKB phenotype or disease signatures.

#### Case 2: Usher syndrome

Usher syndrome, the most common cause of combined deafness and blindness, involves multiple genes including *MYO7A, USH2A, CDH23, PCDH15*, and *USH1C* [Weil et al., 1995, Mathur and Yang, 2015]. The IEKB scientific network subgraph for Usher syndrome visualises the complex interplay among these genes, their shared pathway contexts (e.g., stereocilia formation and mechanotransduction), and cross-phenotype associations spanning sensorineural hearing loss, vestibular dysfunction, and inner ear malformation.

## 4 Web Interface

IEKB provides a responsive, open-access web interface at https://earkb.org, organised into eight top-level modules with gene-specific detail pages linked throughout the site (Figure 5).

**Figure 5:**
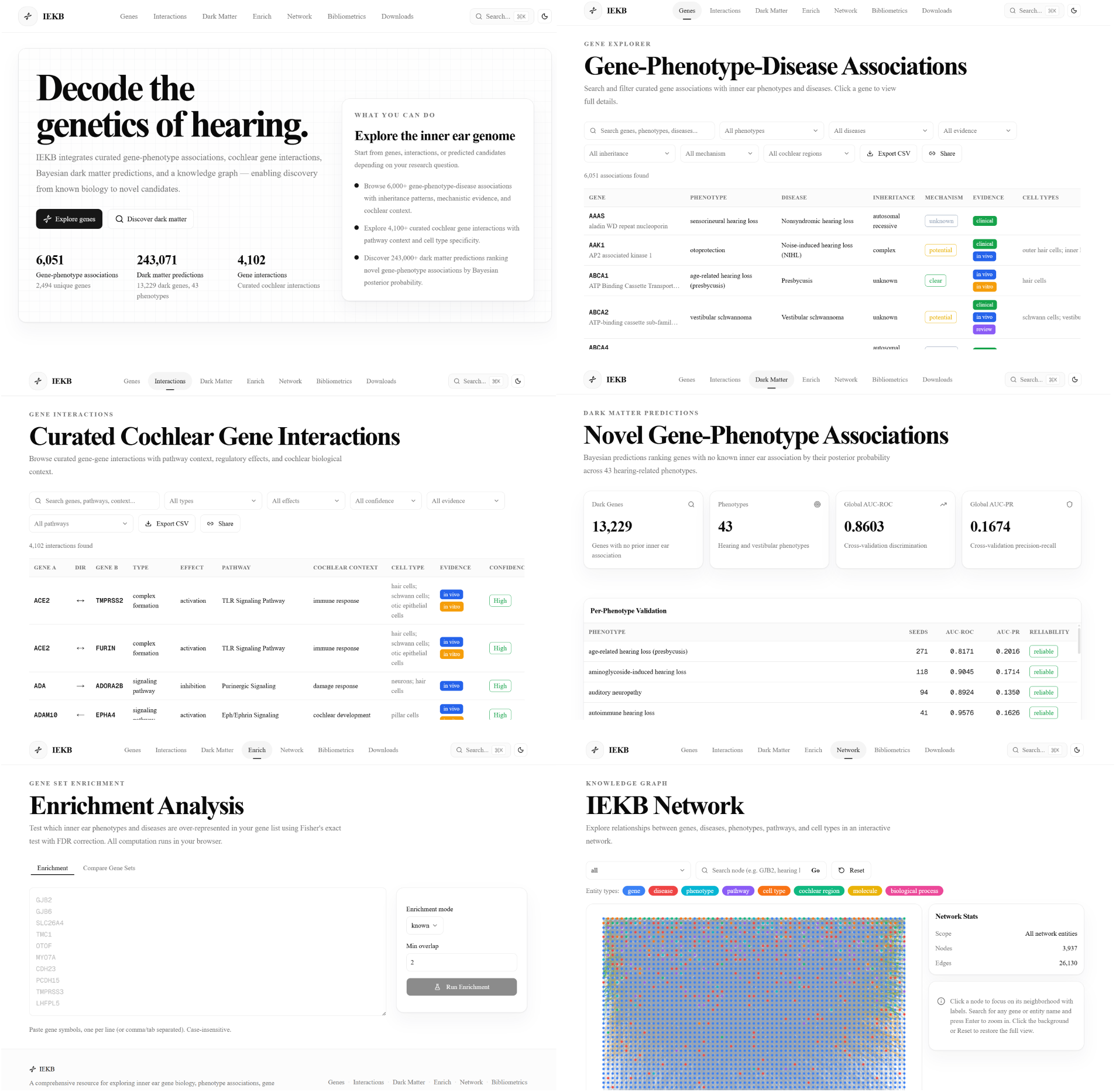
Screenshots of the IEKB web interface. Six representative IEKB interface modules are shown here. *Top left*: Home page displaying summary statistics across the major IEKB data layers and providing quick-start navigation. *Top right*: Gene Explorer page with multi-dimensional filtering (phenotype, disease, inheritance, mechanism, evidence, cochlear region) and paginated results table. *Middle left*: Interactions module showing curated cochlear gene–gene interactions with pathway context, regulatory effects, cell types, and confidence levels. *Middle right*: Dark Matter module displaying Bayesian predictions with per-phenotype validation metrics (AUC-ROC, AUC-PR, reliability) together with an integrated Dark Relation Browser that surfaces known-gene support rankings for each novel candidate. *Bottom left*: Enrichment Analysis module implementing Fisher’s exact test for IEKB phenotype- and disease-based gene-set over-representation analysis, with support for comparing two gene sets. *Bottom right*: Network module rendering the scientific entity network with entity-type colour coding, node search, and neighbourhood exploration.

The **Home** page presents summary statistics across the major data layers—genes, curated associations, interactions, predictions, and network entities—with animated counters and direct entry points to each module.

The **Gene Explorer** supports full-text search by gene symbol, gene name, phenotype, or disease, combined with multi-dimensional filtering by associated phenotype, disease name, evidence level, inheritance pattern, mechanism clarity, and cochlear region. Results are displayed in a paginated, sortable table with one-click CSV export of the current filtered view.

The **Interactions** module presents all 4,102 cochlear gene–gene interactions, filterable by interaction type, regulatory effect, confidence level, pathway context, and evidence level. Each entry links to the supporting PubMed identifiers.

The **Dark Matter** module enables browsing of all 243,071 Bayesian predictions, with filters for phenotype category, minimum posterior probability threshold, and model reliability rating. A dedicated *Dark Relation Browser* tab within the same module exposes the support rankings: for each dark-matter candidate, users can inspect which known genes provide the strongest phenotype-specific evidence, together with calibrated relation scores, feature breakdowns (direct network link, RWR diffusion, gene-group overlap, shared neighbours, and Adamic–Adar index), and natural-language evidence summaries. The relation browser supports multi-parameter filtering by dark gene, known gene, phenotype, reliability, minimum relation score, and minimum dark posterior, as well as sortable columns, pagination, and CSV export of the filtered view. Users can navigate from any prediction or relation to the corresponding gene detail page.

The **Network** module renders the scientific entity network as an interactive graph powered by Cytoscape.js [Franz et al., 2016]. Entities are colour-coded by type (gene, disease, phenotype, pathway, cell type, cochlear region), and users can search for, scope, and focus on specific nodes with neighbourhood highlighting.

The **Enrichment** module implements Fisher’s exact test [Fisher, 1922] for gene-set enrichment analysis. Users input a list of genes and test for over-representation across IEKB phenotype and disease signatures, either using known curated associations alone or combining them with dark matter predictions above a user-defined posterior threshold. A comparison tab allows side-by-side enrichment of two gene sets.

The **Bibliometrics** module summarises the 250,696-article pre-filter retrieval corpus spanning 1814– 2026 across 16,563 screened genes and visualises post-1970 publication trends, top contributing journals, citation patterns, and high-impact articles.

The **Downloads** page (https://earkb.org/downloads) provides all processed core datasets in multiple formats—CSV, JSON, SQLite, and XLSX—accompanied by a data dictionary defining each field. The complete SQLite database (`iekb.db`) can be downloaded for offline querying with any standard SQLite client.

The **Gene Detail** page aggregates all information for a single gene: phenotype associations, interaction partners, dark matter predictions, dark-relation support links (showing known genes that support the candidate or that the gene itself supports), external database links (ClinGen, HHL, OMIM), and supporting literature (Figure 6). A dedicated *Dark Matter Gene* page complements this view by grouping dark-relation evidence by phenotype for a selected candidate gene, with ranked known-gene supporters and concise evidence summaries.

**Figure 6:**
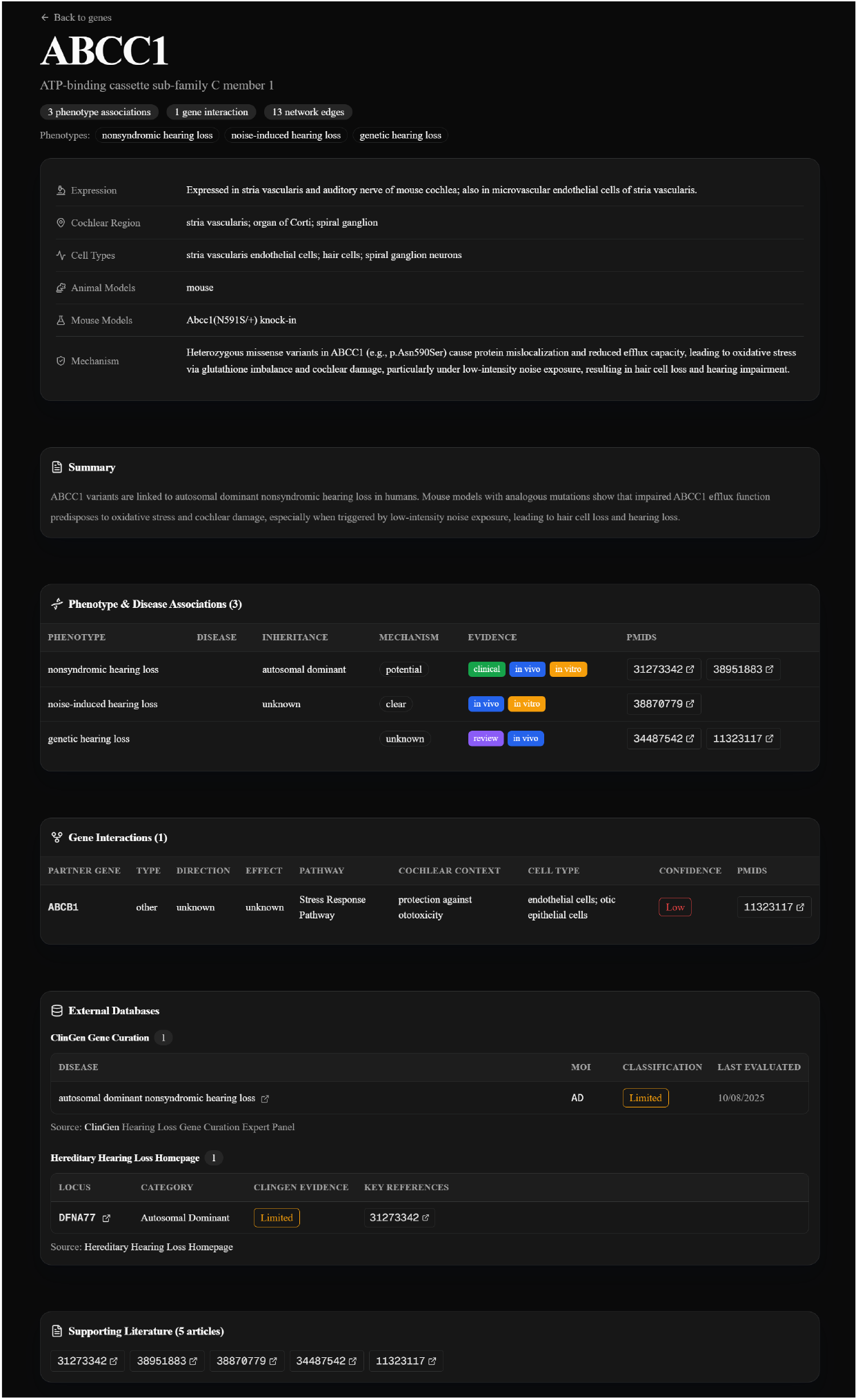
Gene detail page example: *ABCC1*. The gene detail page aggregates all IEKB information for a single gene. For *ABCC1* (ATP-binding cassette transporter C member 1), the page displays: (*i*) gene metadata including expression pattern (stria vascularis, auditory nerve), cochlear regions (stria vascularis, organ of Corti, spiral ganglion), affected cell types (endothelial cells, hair cells, spiral ganglion neurons), animal models (mouse, Abcc1(N591S/+) knock-in), and mechanism description; (*ii*) a narrative summary of the gene’s role in inner ear biology; (*iii*) a table of three phenotype–disease associations (nonsyndromic hearing loss, noise-induced hearing loss, genetic hearing loss) with inheritance, mechanism clarity, and evidence levels; (*iv*) one gene–gene interaction with *ABCB1* in the Stress Response Pathway; (*v*) dark-relation support links showing which known genes provide the strongest evidence for or from this gene; (*vi*) external database cross-references to ClinGen and HHL; and (*vii*) links to five supporting PubMed articles.

Additional features include a global search accessible via keyboard shortcut (Ctrl/Cmd+K), light/dark theme toggle, URL-based sharing of filtered views, and programmatic access to the in-browser database through the browser console (`window.IEKB.query(sql)`). A terms-of-use gate is presented on first visit, informing users of the CC BY 4.0 license and citation requirements.

In addition to the browsing-oriented modules described above, IEKB offers a companion conversational interface, **IEKB QA** (https://qa.earkb.org), which enables researchers to query the knowledge base in natural language and receive structured, evidence-grounded answers. The system employs a multi-phase agent pipeline that classifies incoming questions, retrieves relevant evidence through schema-constrained queries against the IEKB database, ranks candidate genes across multiple scoring dimensions, and synthesises a citation-backed narrative answer streamed to the browser in real time via Server-Sent Events. Users can optionally include dark-matter predictions and dark-relation support evidence, conduct multi-turn follow-up conversations within the same evidence session, and export completed sessions for downstream analysis (Figure 7).

**Figure 7:**
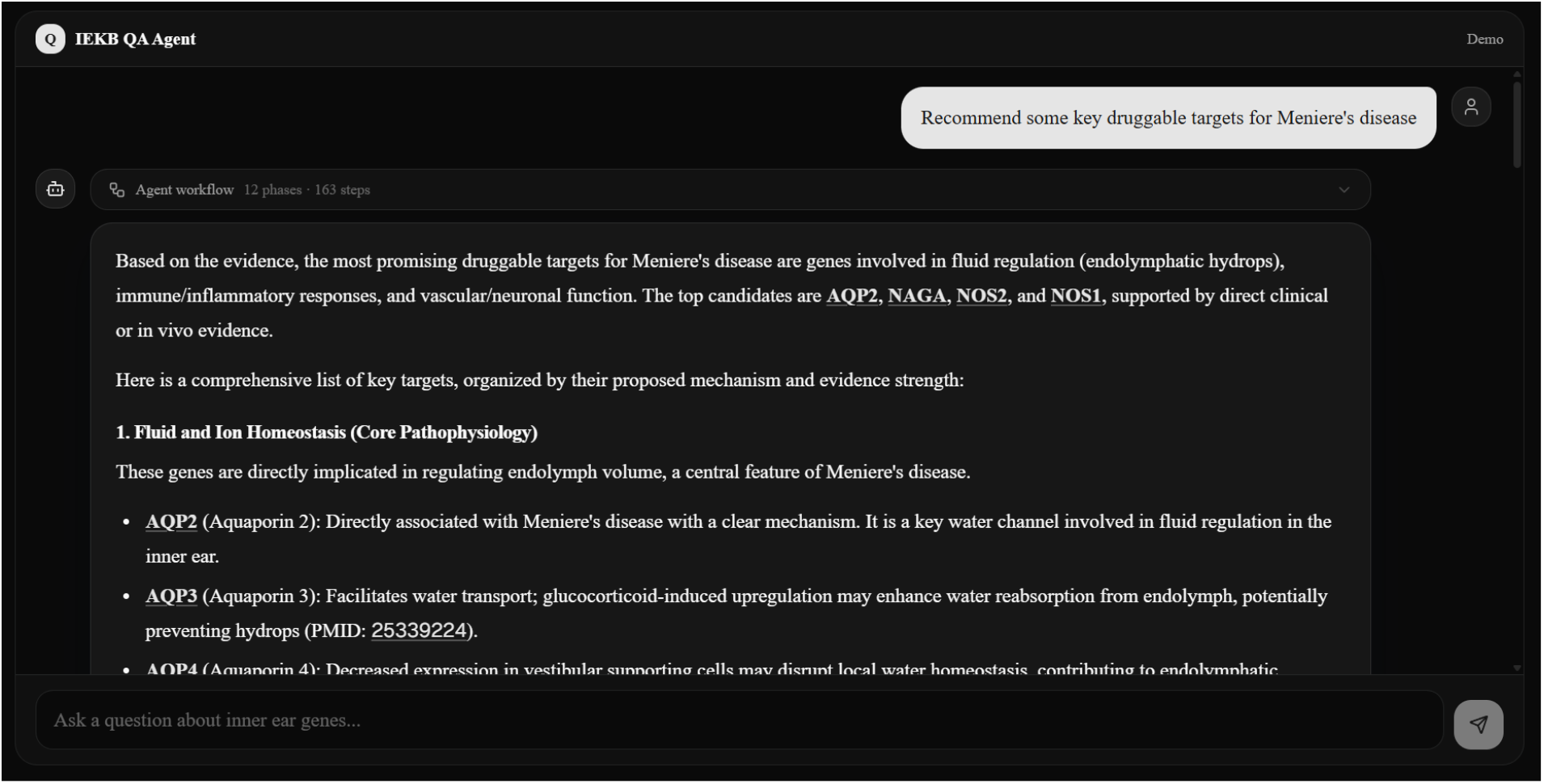
IEKB QA: evidence-grounded question answering interface. The IEKB QA conversational interface (https://qa.earkb.org) allows researchers to pose complex biomedical questions in natural language. The multi-phase agent pipeline classifies the question, retrieves structured evidence from the IEKB database, ranks candidate genes across multiple scoring dimensions, and synthesises a citation-backed narrative answer streamed in real time. *Left*: the chat panel displays agent reasoning steps— question classification, target resolution, evidence retrieval, batch evaluation, and candidate ranking— followed by a streaming narrative answer with inline evidence citations. *Right*: ranked candidate gene cards with multi-dimensional scores, expandable evidence details, and direct links to the corresponding IEKB gene detail pages. Users can toggle dark-matter evidence inclusion, conduct multi-turn follow-up questions, and export completed sessions.

## 5 Database Update

IEKB is designed as a continuously maintained resource rather than a static release. Our update workflow is driven by a customised agent framework that continuously surveys the inner-ear genetics literature for newly published articles linking genes to inner-ear biology, hearing loss, vestibular phenotypes, and related disease mechanisms.

When newly relevant evidence is detected, the system queues the same retrieval, filtering, extraction, normalisation, export, and validation pipeline used for the current release, thereby generating candidate updates to the knowledge base in a timely manner. However, no automatically generated content is published directly. Instead, all newly added or modified records are reviewed by researchers, and each monthly update is released only after manual checking and expert verification of the resulting entries.

During this monthly release cycle, external reference resources—including ClinGen, HHL, and OMIM—are synchronised as needed to preserve alignment with current curation standards. The Bayesian dark matter module and the downstream dark-relation support model are retrained and benchmarked whenever seed sets, interaction evidence, or phenotype definitions change materially; the dark-relation layer is additionally refreshed when the dark-matter candidate list or its posterior thresholds are updated. Any upgrades to the customised agent itself are adopted only after release-level benchmarking confirms that data quality is maintained or improved.

Approved monthly releases are then propagated to the public database and made available through the IEKB website, including the processed download bundles at https://earkb.org/downloads. Users are encouraged to submit corrections, suggestions, or additional data sources through the IEKB website; all user-submitted feedback is evaluated by the curation team before incorporation into subsequent releases.

## 6 Discussion and Future Developments

IEKB addresses a gap not filled by existing hearing-loss resources. HHL, DVD, Gene4HL, SHIELD, and gEAR each contribute valuable locus, variant, or expression information, but none combines curated phenotype–disease evidence, cochlear interactions, Bayesian candidate prioritisation, auditable known-gene support relations for novel candidates, a multi-entity scientific network, bibliometrics, domain-specific enrichment, and downloadable offline data in one openly accessible inner-ear platform (Table 2). This breadth is a central strength of IEKB, allowing users to move from a literature-scale overview to gene-level evidence and network context within a single workflow.

**Table 2:** Comparison of IEKB with representative hearing-loss and inner-ear resources. Feature labels emphasise resource-level functionality rather than implementation details. “–” indicates the feature is not applicable or not provided by the resource.

| Feature | IEKB | Gene4HL | DVD | HHL | SHIELD | gEAR |
| --- | --- | --- | --- | --- | --- | --- |
| Genes / loci covered | 2,494 | 326 | 224 | 158 | Dataset-dependent | Dataset-dependent |
| Phenotype / disease annotations | Extensive | Yes | Variant-linked | Limited | No | No |
| Cochlear gene interactions | Yes | No | No | No | No | No |
| Candidate gene prioritisation | Bayesian | No | No | No | No | No |
| Known-gene support relations | Yes | No | No | No | No | No |
| Scientific entity network | Yes | No | No | No | No | No |
| Domain-specific enrichment | Yes | No | No | No | No | No |
| Bibliometrics | Yes | No | No | No | No | No |
| Downloadable SQLite / offline analysis | Yes | No | No | No | No | No |
| Variant-level data | No | Yes | Yes | No | No | No |
| Expression / omics focus | No | No | No | No | Yes | Yes |
| Open access web resource | Yes | Yes | Yes | Yes | Yes | Yes |

A second strength lies in the curation strategy. Rather than positioning the language model as the product itself, IEKB uses an automated agent-assisted pipeline as an acceleration layer on top of schema constraints, batch-level human monitoring, and domain-expert review. Recent systems such as BiomedCurator and SciDaSynth show the promise of human–AI collaboration for structured literature extraction [Sohrab et al., 2022, Wang et al., 2025], while broader discussions of LLMs in medicine emphasise the importance of careful validation and governance [Thirunavukarasu et al., 2023]. IEKB adopts this conservative stance by keeping expert adjudication in the loop for release-quality data. The dual-tier confidence classification further enables users to distinguish high-confidence, literature-supported entries (Tier 1) from records requiring additional validation (Tier 2), promoting transparent downstream use.

The Bayesian dark matter module introduces a predictive dimension absent from existing inner-ear resources. By combining local network-neighbour support, interaction-network propagation, and gene-group co-occurrence into a unified posterior score, the model can prioritise genes that are plausible from a systems perspective even when direct inner-ear evidence remains sparse. The complementary dark-relation layer goes a step further: rather than presenting a single opaque posterior, it surfaces the specific known genes that most strongly support each dark-matter candidate within a given phenotype context, together with a feature-level decomposition and natural-language evidence summary. This transparency is valuable for experimental follow-up, because investigators can evaluate whether the supporting genes and their evidential basis align with their domain expertise before committing resources. Together with the scientific entity network, these predictive and explanatory layers turn IEKB from a passive catalogue into a hypothesis-generation platform.

The web implementation further strengthens practical utility. Browser-side querying via sql.js [sql.js contributors, 2024], downloadable SQLite assets, bulk exports, and domain-specific enrichment against IEKB phenotype and disease signatures make the resource useful not only for browsing but also for reproducible downstream analysis. Processed release files are openly available from the IEKB downloads portal at https://earkb.org/downloads, enabling direct reuse of curated outputs outside the web interface. The companion IEKB QA interface further complements this browsing-oriented access by allowing researchers to pose complex questions in natural language and receive structured, evidence-grounded answers drawn from the same database, bridging the gap between manual exploration and targeted inquiry. The combination of online exploration, conversational retrieval, and offline reuse is particularly valuable in a field where investigators often move between candidate-gene follow-up, literature review, and comparative interpretation.

Several limitations should be acknowledged. First, literature coverage is constrained by article availability; evidence behind paywalls or absent from PMC/open-access retrieval may be underrepresented. Second, despite human monitoring and expert review, automated extraction can still miss nuance in mechanisms, disease boundaries, or negative evidence. Third, expert curation inevitably involves judgement; inter-annotator agreement metrics quantify part of this variability, and disputed entries are resolved through multi-expert consensus. Fourth, the current release is gene-centred rather than variant-centred, so resources such as DVD [Azaiez et al., 2018] and Gene4HL [Huang et al., 2021] remain complementary for variant interpretation. Fifth, the dark-relation rankings inherit the biases of the underlying STRING network and curated seed sets; relations involving poorly annotated or highly connected hub genes should therefore be interpreted with caution. Sixth, overlap statistics with external resources should be interpreted as release-specific snapshots because reference databases continue to evolve. Seventh, while IEKB QA grounds its answers in structured database evidence, the natural-language synthesis is produced by a large language model, and users should verify that retrieved evidence and narrative phrasing align with the source data before drawing conclusions.

Future development will focus on several directions: (i) deeper full-text processing and release versioning; (ii) additional benchmarking and calibration of candidate prioritisation and dark-relation support models, including incorporation of additional link-prediction features and phenotype-stratified recalibration; (iii) expansion of species coverage beyond the current human-, mouse-, rat-, and guinea-pig-centred evidence base toward broader mammalian representation, including non-human primates; (iv) integration of authoritative single-cell atlas resources to enrich the database with finer-grained gene-expression context; (v) continued expansion of variant-level evidence, the scientific entity network, and domain-specific enrichment modules; and (vi) community-informed curation workflows that scale expert review while preserving traceability and quality control.

## Acknowledgements

We thank all developers and maintainers of the public databases and tools used in this study.

## Funding

Funding details will be finalised in the submission version.

## Conflict of Interest

None declared.

## Data Availability

IEKB is freely available at https://earkb.org. All processed release datasets can be accessed directly at https://earkb.org/downloads without registration or login in CSV, JSON, SQLite, and XLSX formats. The complete SQLite database (`iekb.db`) supports offline querying with any standard SQLite client. The evidence-grounded question-answering interface, IEKB QA, is available at https://qa.earkb.org. All public data are released under the Creative Commons Attribution 4.0 International (CC BY 4.0) license.

## Notes

https://earkb.org

https://qa.earkb.org

### Competing Interest Statement

The authors have declared no competing interest.

## References

Lada A Adamic and Eytan Adar. Friends and neighbors on the web. Soc Netw, 25(3):211–230, 2003. doi: 10.1016/S0378-8733(03)00009-1.

H Azaiez, K T Booth, S S Ephraim, B Crone, E A Black-Ziegelbein, R J Marini, A E Shearer, C M Sloan-Heggen, D Kolbe, T Casavant, et al. Genomic landscape and mutational signatures of deafness-associated genes. Am J Hum Genet, 103:484–497, 2018. doi: 10.1016/j.ajhg.2018.08.006.

Yoav Benjamini and Yosef Hochberg. Controlling the false discovery rate: a practical and powerful approach to multiple testing. J R Stat Soc Series B Stat Methodol, 57:289–300, 1995.

Bryony Braschi, Paul Denny, Kristian Gray, Tamsin Jones, Ruth Seal, Susan Tweedie, Bethan Yates, and Elspeth Bruford. Genenames.org: the HGNC and VGNC resources in 2019. Nucleic Acids Res, 47 (D1):D786–D792, 2019. doi: 10.1093/nar/gky930.

Raquel Cabanillas, Marta Dineiro, Gloria Alvarez Cifuentes, David Castillo, Patricia C Pruneda, Ruben Alvarez, Nuria Sanchez-Duran, Lidia Caixa, Anna Gutiérrez-Arumi, Faustino Ruiz-Juarez, et al. Comprehensive genomic diagnosis of non-syndromic and syndromic hereditary hearing loss in Spanish patients. BMC Med Genomics, 11:58, 2018. doi: 10.1186/s12920-018-0375-5.

Samuel Colvin. Pydantic: data validation using Python type hints. https://docs.pydantic.dev, 2024.

Marina T DiStefano, Sarah E Hemphill, Andrea M Oza, Rebecca K Siegert, Andrew R Grant, Madeline Y Hughes, Sami S Amr, and ClinGen Hearing Loss Clinical Domain Working Group. ClinGen expert clinical validity curation of 164 hearing loss gene-disease pairs. Genet Med, 21(10):2239–2247, 2019. doi: 10.1038/s41436-019-0487-0.

Ronald A Fisher. On the interpretation of χ2 from contingency tables, and the calculation of P. J R Stat Soc, 85:87–94, 1922.

Max Franz, Christian T Lopes, Gerardo Huck, Yue Dong, Onur Sumer, and Gary D Bader. Cytoscape.js: a graph theory library for visualisation and analysis. Bioinformatics, 32:309–311, 2016. doi: 10.1093/bioinformatics/btv557.

GBD 2019 Hearing Loss Collaborators. Hearing loss prevalence and years lived with disability, 1990–2019: findings from the Global Burden of Disease Study 2019. Lancet, 397:996–1009, 2021. doi: 10.1016/S0140-6736(21)00516-X.

Shasha Huang, Guihu Zhao, Jie Wu, Kuokuo Li, Qiuquan Wang, Ying Fu, Honglei Zhang, Qingling Bi, Xiaohong Li, Weiqian Wang, et al. Gene4HL: an integrated genetic database for hearing loss. Front Genet, 12:773009, 2021. doi: 10.3389/fgene.2021.773009.

David P Kelsell, Jonathan Dunlop, Holger P Stevens, Nicholas J Lench, Jia N Liang, Gareth Parry, Robert F Mueller, and Irene M Leigh. Connexin 26 mutations in hereditary non-syndromic sensorineural deafness. Nature, 387:80–83, 1997. doi: 10.1038/387080a0.

Prem Mathur and Jun Yang. Usher syndrome: hearing loss, retinal degeneration and associated abnormalities. Biochim Biophys Acta, 1852:406–420, 2015. doi: 10.1016/j.bbadis.2014.11.020.

Cynthia C Morton and Walter E Nance. Newborn hearing screening — a silent revolution. N Engl J Med, 354:2151–2164, 2006. doi: 10.1056/NEJMra050700.

Yuji Nakajima. Signaling regulating inner ear development: Cell fate determination, patterning, morphogenesis, and defects. Congenit Anom (Kyoto), 55(1):17–25, 2015. doi: 10.1111/cga.12072.

Daniel N Nicholson and Casey S Greene. Constructing knowledge graphs and their biomedical applications. Comput Struct Biotechnol J, 18:1414–1428, 2020. doi: 10.1016/j.csbj.2020.05.017.

Joshua Orvis, Brian Gottfried, Jayaram Kancherla, Ronna S Adkins, Yang Song, Amiel A Dror, David Olley, Kevin Rose, Erika Chrysostomou, Michael C Kelly, et al. gEAR: gene expression analysis resource portal for community-driven, multi-omic data exploration. Nat Methods, 18:843–844, 2021. doi: 10.1038/s41592-021-01200-9.

Heather Piwowar, Jason Priem, Vincent Lariviere, Juan Pablo Alperin, Lisa Matthias, Bree Norlander, Ashley Farley, Jevin West, and Stefanie Haustein. The state of OA: a large-scale analysis of the prevalence and impact of open access articles. PeerJ, 6:e4375, 2018. doi: 10.7717/peerj.4375.

Eric W Sayers, Evan E Bolton, J Rodney Brister, Kathi Canese, Jessica Chan, Donald C Comeau, Ryan Connor, Kathryn Funk, Chris Kelly, Sunghwan Kim, et al. Database resources of the National Center for Biotechnology Information. Nucleic Acids Res, 50(D1):D13–D25, 2022. doi: 10.1093/nar/gkab1112.

A Eliot Shearer, Michael S Hildebrand, Amanda M Odell, and Richard J. H. Smith. Genetic hearing loss overview. GeneReviews® [Internet]. University of Washington, Seattle; NCBI Bookshelf, 2025. Last revision 3 April 2025. https://www.ncbi.nlm.nih.gov/books/NBK1434/.

Jing Shen, Deborah I Scheffer, Kelvin Y Kwan, and David P Corey. SHIELD: an integrative gene expression database for inner ear research. Database (Oxford), 2015:bav071, 2015. doi: 10.1093/database/bav071.

Mohammad Golam Sohrab, Khoa N. A. Duong, Masami Ikeda, Goran Topic, Yayoi Natsume-Kitatani, Masakata Kuroda, Mari Nogami Itoh, and Hiroya Takamura. BiomedCurator: Data curation for biomedical literature. In Proceedings of the 2nd Conference of the Asia-Pacific Chapter of the Association for Computational Linguistics and the 12th International Joint Conference on Natural Language Processing: System Demonstrations, pages 63–71, Taipei, Taiwan, 2022. Association for Computational Linguistics. doi: 10.18653/v1/2022.aacl-demo.8. URL https://aclanthology.org/2022.aacl-demo.8/.

sql.js contributors. sql.js – SQLite compiled to JavaScript. https://sql.js.org, 2024.

Arun James Thirunavukarasu, Daniel Shu Wei Ting, Karthik Elangovan, Laura Gutierrez, Ting Fang Tan, and Daniel S W Ting. Large language models in medicine. Nat Med, 29:1930–1940, 2023. doi: 10.1038/s41591-023-02448-8.

Hanghang Tong, Christos Faloutsos, and Jia-Yu Pan. Random walk with restart: fast solutions and applications. Knowl Inf Syst, 14:327–346, 2008. doi: 10.1007/s10115-007-0094-2.

Giovanni Visonà, Emmanuelle Bouzigon, Florence Demenais, and Gabriele Schweikert. Network propagation for GWAS analysis: a practical guide to leveraging molecular networks for disease gene discovery. Brief Bioinform, 25(2):bbae014, 2024. doi: 10.1093/bib/bbae014.

Daniel Walls, Hela Azaiez, and Richard J. H. Smith. Hereditary hearing loss homepage. https://hereditaryhearingloss.org/, 2026. Accessed 22 March 2026.

Xingbo Wang, Samantha L Huey, Rui Sheng, Saurabh Mehta, and Fei Wang. SciDaSynth: Interactive structured data extraction from scientific literature with large language model. Campbell Syst Rev, 21 (4):e70073, 2025. doi: 10.1002/cl2.70073.

D Weil, S Blanchard, J Kaplan, P Guilford, F Gibson, J Walsh, P Mburu, A Varela, J Levilliers, M D Weston, et al. Defective myosin VIIA gene responsible for Usher syndrome type 1B. Nature, 374: 60–61, 1995. doi: 10.1038/374060a0.

